# The action of antimicrobial peptide LL37 is slow but effective against non-growing *Escherichia coli* cells

**DOI:** 10.1101/2022.08.29.505736

**Authors:** Salimeh Mohammadi, Sattar Taheri-Araghi

**Affiliations:** Department of Physics and Astronomy, California State University, Northridge, CA 91330, USA

## Abstract

Antimicrobial peptides (AMPs) are amino-acid based antibiotics that primarily attack bacteria by perturbing their membranes. It has long been suggested that AMPs are effective against non-growing cells that tolerate conventional antibiotics. Despite recent advances in quantifying the action of AMPs at the single-cell level, we still do not have a clear picture of how this action is correlated with the physiology of target cells. Here we take complementary approaches, from single-cell and population-level experiments, to investigate the efficacy of human AMP LL37 against *Escherichia coli* cells at different growth phases. We first analyze time-lapse, single-cell data of the action of LL37 peptides on exponentially growing cells, which reveals that they act *faster* on long, dividing cells than on small, newborn cells. Next, we test the consequence of this cell-age dependence on the efficacy of LL37 against non-growing *E. coli* cells in stationary phase. We observe a consistent trend that the action of LL37 is, on average, ≈5.0 minutes slower on non-growing cells as compared to that on exponentially growing cells. However, this difference in the rate of action is not reflected in the minimum bactericidal concentration (MBC) of LL37 peptides. Contrary to our expectation, the MBC measured for non-growing cells is smaller than that for exponentially growing cells, indicating that over a long period of time the LL37 peptides are more potent against non-growing cells.

**Author summary:** Antibiotic treatments can fail due to regrowth of a bacterial subpopulation that start proliferation after the treatment is over. The regrowth is often from non-growing, dormant cells that persist the action of antibiotics without being resistant. In this work, we demonstrate that human antimicrobial peptide LL37 is potent against non-growing *Escherichia coli* cells.

## Introduction

Antibiotic therapies can fail due to regrowth of persister cells that start proliferation after the antibiotic treatment is removed [1–4]. Persister cells are often observed in the form of non-growing (dormant) cells that survive therapy without developing genetic resistance [1–3, 5–7]. This phenotypic heterogeneity is common when an antibiotic targets pathways related to bacterial growth and cell-cycle progression [1–3,5,6]. This is, thus, of fundamental importance in science and medicine to identify antibiotic agents that combat non-growing cells. Antimicrobial peptides (AMPs) are one of the candidates that have long been suggested to be effective against dormant cells [8,9].

AMPs attack bacteria first by electrostatic absorption onto their membrane and then by hydrophilic insertion into their lipid bilayers [8–12]. Most AMPs have amphiphilic structures which incorporate cationic side chains that bind to negatively charged membranes and non-polar side chains that facilitate penetration into lipid bilayers [8,10–15]. Even though we have recently observed that AMPs interactions go beyond membrane perturbations, the membrane interaction is indeed the initial step of the bacterial killing process [16–18].

To date, we do not fully understand how the action of AMPs is affected by the physiology of target cells. Specifically, the correlation between the action and the physiological parameters that define progression of the cell-cycle, cell age, or cell size is not well-quantified. The first observation of the cell-cycle dependence of the action of AMPs is from the work of Weisshaar lab that demonstrated LL37 peptides first bind to the septum area in *Escherichia coli* cells, leading to a higher activity of LL37 peptides on dividing cells [19]. This observation raises the question of whether AMPs are potent against cells in stationary phase [20].

In this work, we integrate single-cell and population-level approaches to systematically investigate the efficacy LL37 peptides against non-growing *E. coli* cells. We start by analyzing the cell-size dependence of the action of LL37 peptides, which confirms that the susceptibility of *E. coli* cells increases over the bacterial life cycle. That is, statistically, large cells (near the stage of cell division) are more susceptible to LL37 peptides as compared to the small cells.

Next, we enforce a controlled starvation state on cells by depleting nutrient for a fixed duration of time to study effectivity of LL37 peptides on non-growing cells. In the starvation state, where the frequency of cell division is reduced to zero, the rate of action of LL37 peptides becomes slow, lagging behind that in the exponentially growing cells.

However, this reduction in the rate of action is not directly reflected in the minimum bactericidal concentration (MBC) measurements. Surprisingly, we find that the MBC of LL37 peptides is smaller in starving cultures. This indicates that, despite the slower action, LL37 peptides are more potent against non-growing cells than on exponentially growing cells. This, indeed, is a favorable observation in terms of the potential medical application of AMPs as antibiotic agents.

## Materials and methods

### Strain and growth condition

In all experiments we used a derivative of an *Escherichia coli* K12 strain, NCM3722, that was constructed and tested by Kustu and Jun labs [21,22]. We used ST08, a non-motile derivative of NCM3722 (ΔmotA). The growth media that we used is a MOPS based rich defined media (RDM) developed by Fred Neidhardt [23], which is commercially available from Teknova Inc. In this media, the average doubling time of ST08 strain is about 23 minutes at 37 °C.

### Sample preparation

The cell culture was carried out in multiples stages: (1) the seed culture; (2) pre-culture; (3) experimental culture (in exponential phase); and (4) starved culture (if applicable).

(1) The seed culture was inoculated from an isolated colony on a Lysogeny Broth (LB) agar plate into 3mL of RDM in a culture tube (950 mm × 150 mm) incubated at 37 °C in a water bath shaker. The colonies on the LB agar plate were prepared by streaking a −80 °C glycerol stock on the plate, first incubated at 37 °C for 16-24 hours and then kept in the 4 °C fridge.

(2) For the pre-culture, the seed culture was diluted 1000-fold in the identical growth medium (RDM) after overnight growth and incubated in the 37 °C water bath shaker until it reached mid-exponential phase.

(3) For the experiments with exponentially growing cells, the cells were harvested at the optical density of 0.2 (measured with light at 600 nm, referred to as OD_600_) and then diluted to OD_600_ of 0.002 for the microscopy and MBC measurement experiments.

(4) For the experiments with starved cells, 1 mL of the culture was concentrated and washed twice in a micro-centrifuge tube with the no-nutrient buffer. Next, the culture was diluted to OD_600_ =0.002 to avoid any potential inoculum effect [24–27]. The culture was then incubated for three hours for transition to the stationary phase.

### Antimicrobial peptide

We used antimicrobial peptide LL37 (AnaSpec, California). The product was shipped in the form of dry powder in vials and 400 M stocks were prepared in autoclaved double-distilled water. The liquid stock was stored at −20 °C. The net peptide content of the product was measured to be 75% by elemental analysis of the carbon, hydrogen, and nitrogen (CHN analysis) conducted by the manufacturer. Special handling precautions were used for any solutions containing antimicrobial peptides: low protein binding supplies including pipette tips (Low Retention Aerosol Barrier, FisherBrand), micro-centrifuge tubes (Protein LoBind, Eppendurf), and microplates (Ultra-Low attachment, Corning) were used to avoid attachment of AMPs to the experimental supplies.

### Live cell microscopy

The microscopy procedure in this work is similar to that used in reference [24]. Individual *E. coli* cells were immobilized by agarose pads (5% agarose gel) for live microscopy to monitor the activity of LL37 peptides on them. Inspired by others [28, 29], we used patterned agarose gel in a housing for long term microscopy experiments. The patterns provide parallel channels to spread the cells under the agarose pad. The housing reduces the evaporation, thus helping long term (few hours) microscopy at 37 °C. For each experiment, several fields of view were studied from the same sample. The interval between the time-lapse images is 1 minutes for all the experiments.

### Live/Dead^®^ stain

To differentiate between live and dead *E. coli* cells in the single-cell microscopy experiments we used Live/dead^®^ BacLight™ fluorescent stain kit from Invitrogen, California. This kit includes two nucleic acid stains: propidium iodide (PI) and SYTO9. PI is the dead stain that can enter into the cytoplasm of the cells with perturbed membrane and fluoresce once bound to the DNA, with the excitation/emission wavelengths of 480/500 nm. The SYTO9 is the live stain that can cross all bacterial membrane, including those with intact membranes, and fluoresce with the excitation/emission wavelengths of 490/635 nm.

To use the kit, we first incubated 1mL of the culture with a total of 3 μL of 1:1 mixture of the dyes. After 5 minutes of incubation, the culture was transferred under agarose gel pad for time-lapse microscopy. The agarose gel pad contained a lethal dosage (5 μM) of LL37 peptides and a 1:1 mixture of both components of the kit.

### Determination of the minimum bactericidal concentration of LL37 peptides

To determine the minimum bactericidal concentration (MBC) of LL37 peptides in an *E. coli* culture, we monitored the viability of cells on LB-agar plates from cultures treated with varying dosages of LL37 peptides. The culture was prepared as detailed in the Sample Preparation section, dilute to OD_600_=0.002, and then treated with different concentrations of LL37 peptides ranging from 0.25 to 2.00 μM in a 96-well plate (Ultra-Low attachment, Corning), which was incubated on a shaker at 37°C for two hours. Next, 5 μL of culture from each well (bearing varying dosages of LL37 peptides) was transferred to an LB-agar plate. The growth on LB-agar plate is detectable after 16 to 24 hours of incubation at 37 °C. The lowest concentration of LL37 peptides leading to a complete growth inhibition on the LB plate was considered as the MBC.

## Results

### Susceptibility of *E. coli* cells to LL37 peptides increases over cells’ life cycle

To monitor the inhibition of bacterial growth at single-cell resolution and to quantify the distribution/penetration of AMPs in target cells, we previously performed live microscopy on *E. coli* cells using a dye-tagged version of LL37 peptide, 5-FAM-LC-LL37 (Anaspec, California) [24]. We harvested cells from a culture in mid-exponential phase and brought them under a live microscopy setting, where they were immobilized under an agarose gel pad containing the growth media (RDM) and a lethal concentration (10 μM) of 5-FAM-LC-LL37 peptides. The time-lapse data from the phase contrast and fluorescent microscopy showed that upon cell death, a large number of peptides was absorbed and trapped in the cytoplasmic area of the cell [24].

Here we analyze the growth trajectory of each individual cell prior to cell death to examine the correlation between the cell-size and the time of cell death (Fig. 1)^1^. To visualize this correlation, we use a color-coded plot in Fig. 1A, which reflects the size evolution based on the cell-length at the beginning of the experiment: blue shades correspond to the small cells and red shades correspond to the large cells. The histogram on the bottom of Fig. 1A depicts the probability distribution of death time and the horizontal bar depicts the gradient of the death time, color-coded based on the average initial cell-size. The histogram and the color gradient of the horizontal bar indicate that there is a negative correlation between cell-length and the timing of cell death: the large cells die first, small cells next. Fig. 1C directly demonstrates this negative correlation between the susceptibility and cell-size.

**Fig 1.**
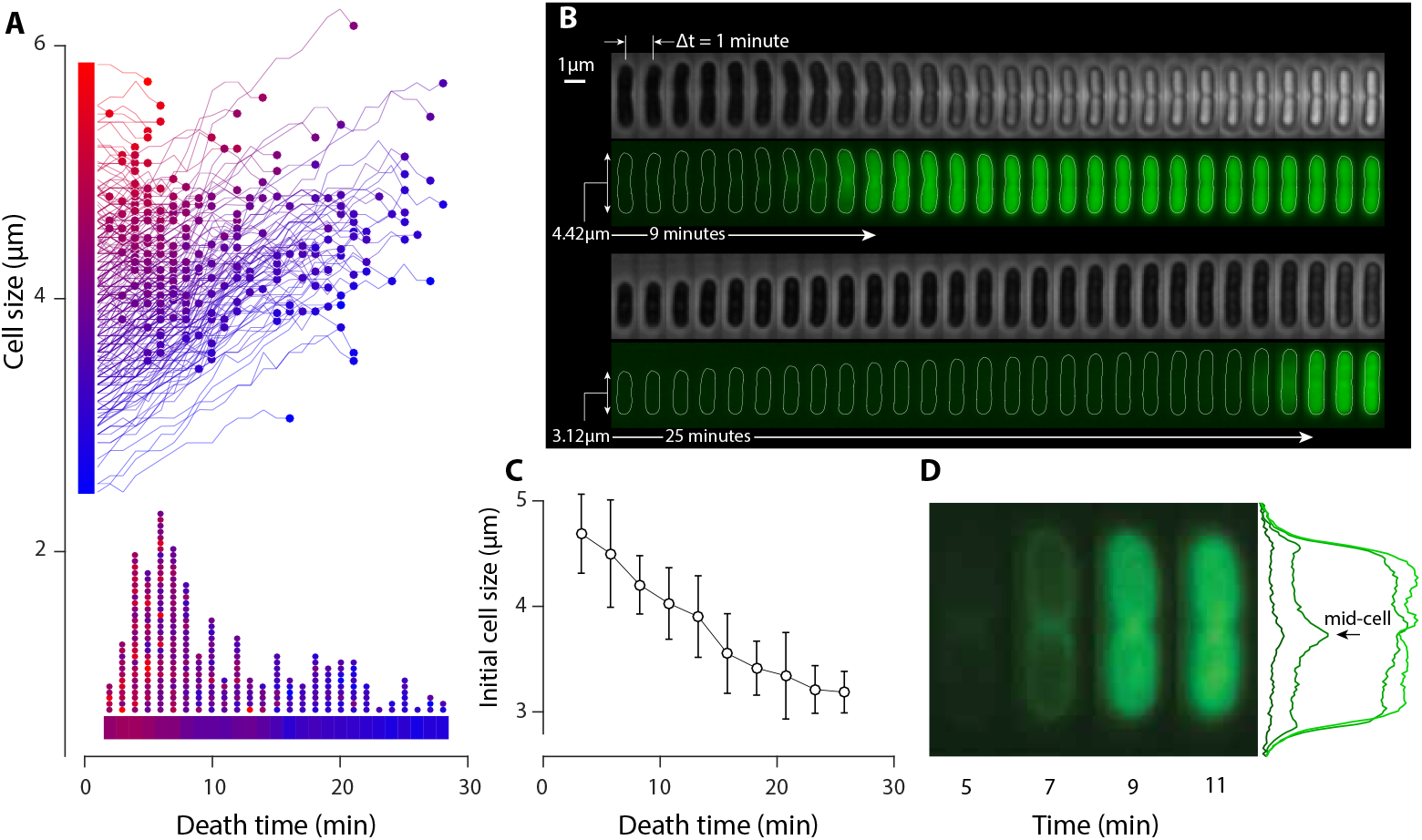
The single-cell evidence of the cell-size dependence of LL37 action. (A) Growth of individual *E. coli* cells prior to cell death by a lethal dosage (10 μM) of 5-FAM-LC-LL37 peptides. Curves are color-coded based on the initial cell size: the blue to red gradient denotes small to large as indicated by the vertical bar next to the y-axis. The circle at the end of each curve depicts the time of cell death. The histogram of death time at the bottom of the panel shows small cells (blue) die later than large cells (red). (B) Sample phase-contrast and fluorescent images of a large and a small *E. coli* cell. Death is accompanied by a rapid translocation of peptides (green signal) into the cells. The large cell dies after 9 min while the small cell dies after 25 min. (C) The negative correlation of initial cell-size and death time shows that large cells are more susceptible to LL37 than small cells. (D) The fluorescent intensity profile of 5-FAM-LC-LL37 shows that peptides preferentially bind around the septum of dividing cells. The curves with increasing intensity correspond to the four snapshots from left to right, respectively.

After cell death, the cell-size distribution does not show substantial correlation with the timing of the death or with the initial cell size (Fig. S1). Fig. 1B shows samples of one small (3.12 μm) and one large (4.42 μm) cell where the timings of cell death (AMP translocation into the cytoplasmic area) are different (9 and 25 minutes) but the cell-size after death is relatively similar. This indicates that the small cell continued to grow until it became susceptible to LL37 peptides.

### The LL37 peptides mostly accumulate around the septum position prior to cell death

The fluorescent images reveal the absorption and accumulation of dye tagged peptides (5-FAM-LC-LL37) on the membrane and into the cytoplasmic area of *E. coli* cells. Consistent with the previous work from Weisshaar lab, the peptides first bind to and accumulate around the septum of dividing cells [19]. We do not understand mechanistically how peptides identify the septum, but one plausible explanation is the tendency of LL37 peptides to preferentially bind to negative Gaussian curvatures (saddle-point curvatures) as shown by Wong lab [30,31]. The only surface on *E. coli* membrane with negative curvature is the septum where the cell-width is narrow in the middle of the cell. In many cases, a bulging and a rupture occurring at the septum position is observed in phase contrast and fluorescent images (Video. S1).

### Absence of nutrient can temporarily stop bacterial growth without compromising cell viability

The observation that susceptibility is correlated with cell division raises the question of whether the efficacy of LL37 peptide is influenced by sub-optimal physiological conditions where the frequency of cell division is reduced. To answer this question, we created and tested a controlled starvation condition for *E. coli* cells with the goal of effectively suppressing growth and cell division without compromising cell viability. To this end, we used the buffer component of the growth media (RDM) with no added sugar or supplements. Cells inoculated in this buffer did not show any population growth detectable by spectrophotometer (Genesys 2000, Fisher Scientific).

To monitor the transition of cells from exponential phase to this buffer, we utilized the microfluidic “mother machine,” which allows for continues and rapid environmental control as well as microscopy at single-cell resolution. We monitored around 400 individual cells over tens of generations as they experience the transition from RDM to buffer and back to RDM. Specifically, we first grew cells in RDM for about ten hours in mother machine and then switched to the buffer for three hours. The fast infusion ensures rapid transition of the environment to the non-nutrient buffer. We observed that after the media is switched to the buffer, the growth rate (i.e., the elongation rate) of individual cells rapidly drops to zero and the division rate decreases after about a 60-minute delay (Fig. 2AB). Over three hours in the buffer, the growth was largely paused. Lastly, we switched again back to RDM to evaluate viability of the cell. And we evidenced that all cells started growing again soon after the infusion of RDM and the rate of cell division was recovered shortly after that.

**Fig 2.**
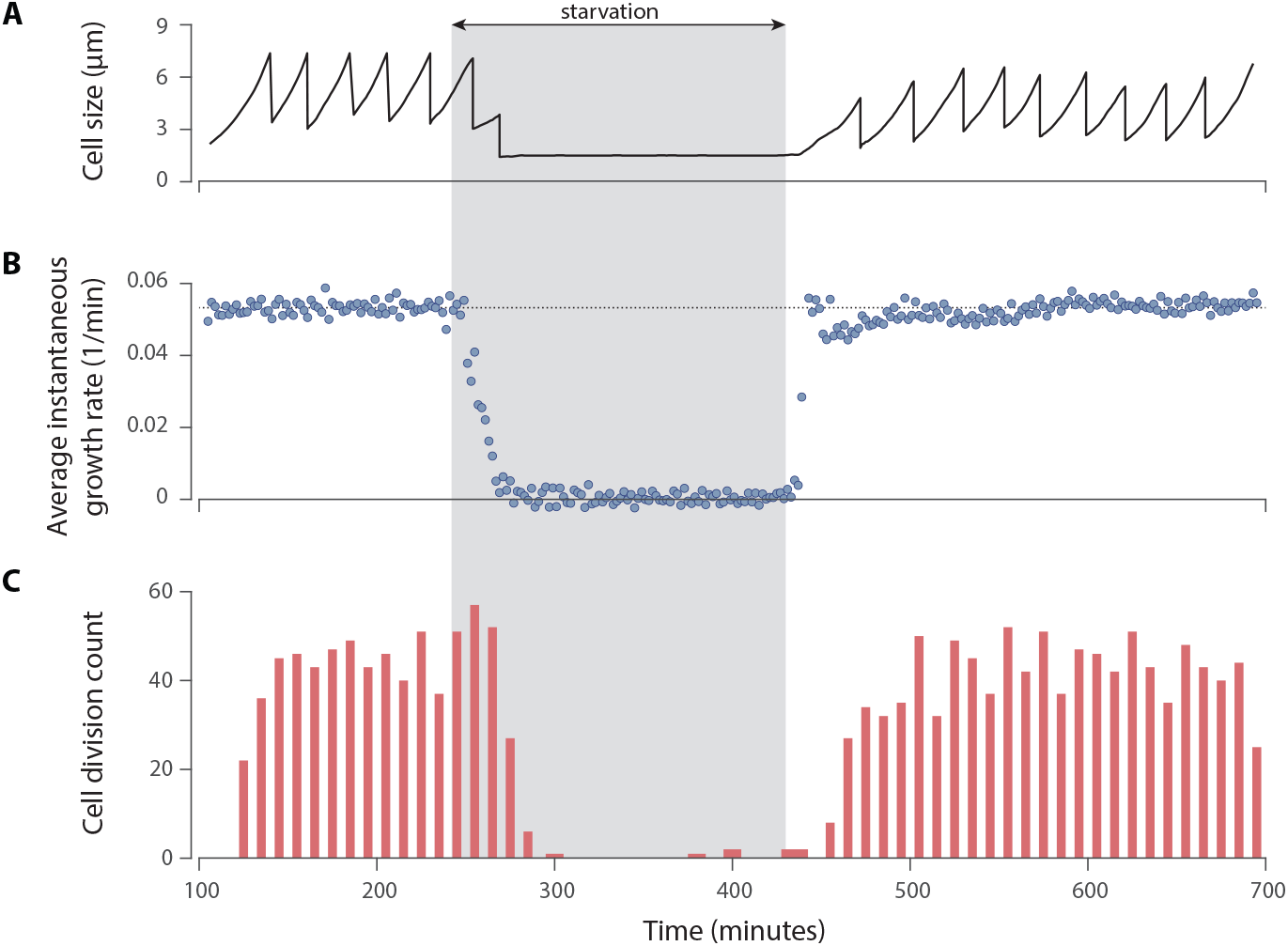
Growth inhibition and viability of *E. coli* cells in a no-nutrient buffer. (A) Growth and division of a sample cell shows that growth is paused after two cell divisions during the starvation period. But growth is rapidly recovered after the growth media is restored in the environment. (B) The instantaneous growth rate (elongation rate) of cells drops to zero during the starvation period and recovers to normal rate once the growth media is replenished. (C) Cell divisions stop after a delay during the starvation period and recover afterwards.

In what follows in this work, we use the transition of growth media to the buffer for three hours as a method to induce stationary phase on *E. coli* cells for investigating the action of LL37 peptides on non-growing cells.

### Single-cell data shows that cells in buffer (in absence of nutrients) tolerate action of LL37 peptides for a longer period of time

To monitor the bactericidal activity of LL37 peptides on individual *E. coli* cells in either stationary or exponential phase we used live/dead^®^ BacLight bacterial viability kit (Life Technologies, California). This allows us to use the un-tagged, original version of LL37 peptides, which is an advantage over the method we presented in Fig. 1^2^.

The kit utilizes a mixture of two nucleic acid stains: (2) propidium iodide (PI), a red-fluorescent dye that penetrates only into bacteria with damaged membrane and stains their DNA; and (2) SYTO9, a green-fluorescent dye that can stain the DNA of live cells with intact membrane. The difference in spectral characteristics and the ability to penetrate live/dead bacteria makes these stains suitable to study temporal dynamics of cell death in time-lapse microscopy experiments.

The disadvantage of using this kit is that the response time of the dyes is slow, thus, not providing us with a sharp, high temporal-precision detection of death time. For this reason, the analysis and the conclusions in this section is based on the average intensity variations and the time shifts that we observe as a function of the cell’s physiological state.

The *E. coli* cells were prepared in either exponential phase or stationary phase (as detailed in Material and Methods section) and were transferred under an agarose pad for microscopy. The pad contained the corresponding media (RDM or buffer), a 5 μM of LL37 peptides, and a mixture of the PI and SYTO9 stains.

Fig. 3A shows a representative time-lapse result of an *E. coli* cell in phase contrast and two fluorescent channels revealing signals from both STYO9 and PI. Since the cell is live at the beginning of the experiment, we have an initial gradual increase in the SYTO9 green signal, which indicates that there is a time-scale for a complete interaction of SYTO9 with live cells. Once the cell is permeabilized by LL37 peptides, the SYTO9 signal starts to fade away and, concurrent with that, the red signal from PI increases.

**Fig 3.**
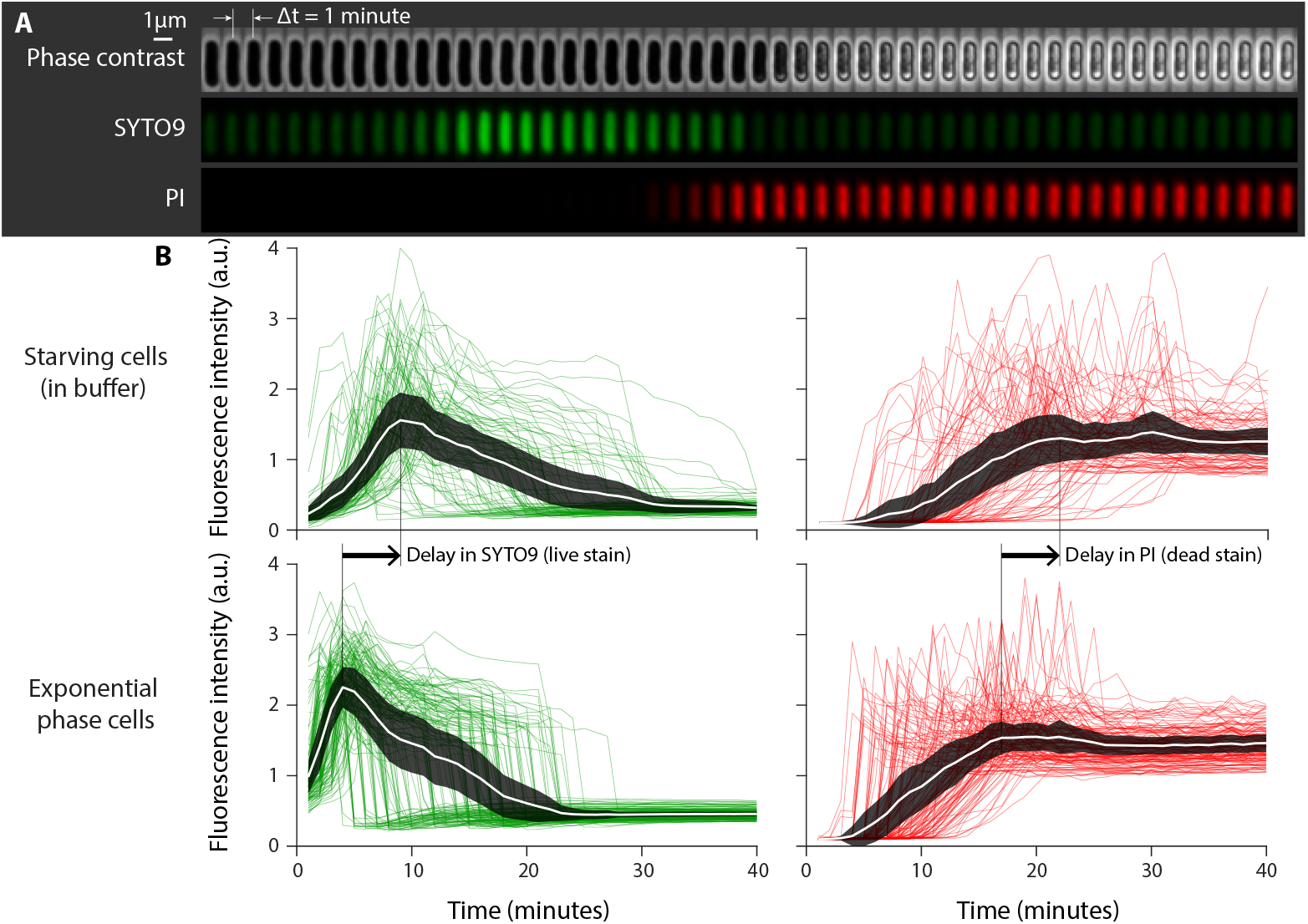
Single-cell microscopy shows the resistance of non-growing cells to LL37 peptides. (A) Sample time-lapse images of an *E. coli* cell under attack by a lethal dosage (5 μM) of LL37 peptide while the cell viability is monitored with live/dead^®^ stains. SYTO9 signal (green) disappears because of cell death while the propidium iodide (PI) signal (red) appears because of increased membrane permeability. (B) Fluorescence intensity profile of SYTO9 (green, left panels) and PI (red, right panels) of 97 starving cells (top panels) and 139 cells from exponential phase culture (bottom panels). The starving cells have been in buffer for three hours prior to the microscopy experiment. Comparison of the peak of SYTO9 signal and the plateau of PI signal shows that death of starving cells occurs after about a 5-minute delay.

Fig. 3B depicts the intensity of the stains’ fluorescent signal (averaged over the cell area) for 139 cells in exponential phase and 97 cells in starvation phase. We observe the same qualitative trend in the variations of the fluorescent signals, indicating that all cells are live at the beginning of the experiment and die during the course of the time-lapse imaging as a results of LL37 peptides.

However, a comparison of the data from exponentially growing cells and stationary phase cells reveals that, despite the same trend, action of LL37 peptides on starving cells lags behind that on exponentially growing cells. The the SYTO 9 signal average as a peak (Fig. 3B left panels), which occurs about 5 minutes later for the cells in starvation state (top left panel). The PI signal average has a plateau, which occurs with the same delay for starving cells (Fig. 3B right panels).

### Minimum bactericidal concentration of LL37 peptides against starving cells

The observation that non-growing cells tolerate LL37 peptides for a longer time period raises the question of how a culture of starving bacteria in stationary phase responds to AMPs. To answer this question, we compared the minimum bactericidal concentration (MBC) of LL37 peptides in cultures of starving cells with that in cultures of exponentially growing cells.

To measure the MBC, we incubated cells with varying dosages of LL37 peptides for two hours and examined their viability on LB-agar plates (for details see Materials and Methods section). We conducted parallel experiments on both exponentially growing cells and starving cells for the sake of comparison and, in both cases, we used the same cell density to avoid any inoculum effect (variation of MBC due to cell density [24–26]).

A typical, general dynamic for the action of antibiotics states that the rate of action is inversely related to the MBC. That is, the faster killing rate of an antibiotic would correspond to a smaller MBC [32].

Interestingly, here we observe an opposite relationship where the MBC of LL37 is smaller when acting on starving *E. coli* cells, though at a slower rate. The MBC data is presented in Fig. 4A, where the MBC on non-growing, starving culture is 0.54 ± 0.12 *μ*M and for exponentially growing cells is 0.88 ± 0.12 *μ*M.

**Fig 4.**
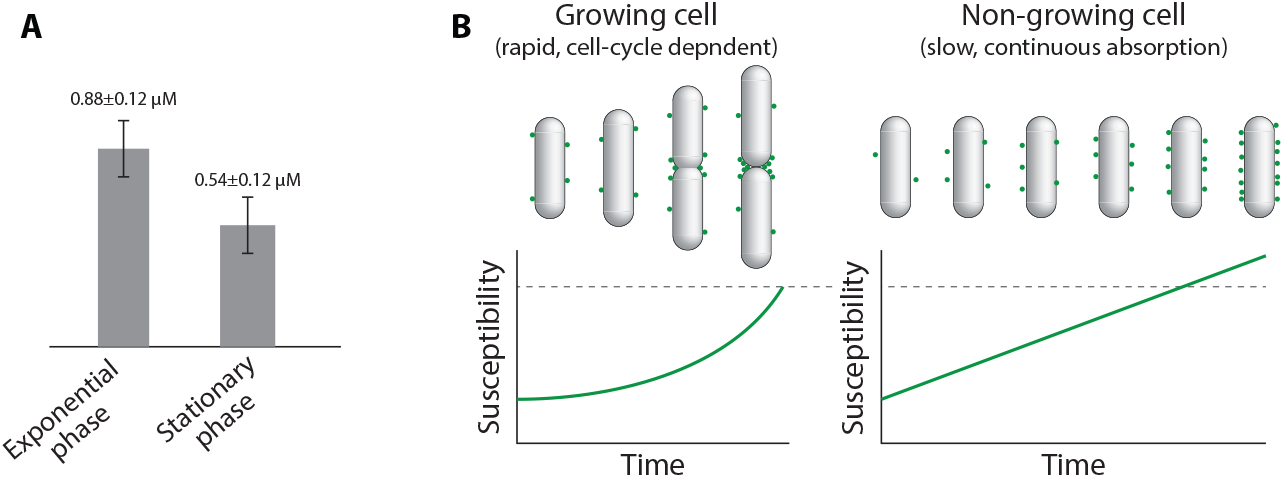
(A) The MBC of LL37 peptides in exponentially growing and in non-growing, starving *E. coli* cultures. (B) A hypothetical model where the susceptibility of growing cells is dependent on cell age. But non-growing cells are susceptible to AMPs due to a slow, yet continuous and long term absorption of peptides on their membranes.

## Discussion and Conclusion

One of the reasons for the failure of antibiotic therapies is the reduced susceptibility of non-growing, dormant cells to antibiotics since many conventional antibiotics target pathways involved in the growth and cell-cycle progression [1, 6, 33–37]. However, antimicrobial peptides have been suggested to retain their potency against non-growing cells as their membrane-perturbation action is, presumably, independent of the bacterial growth state [8, 9].

In this work, we used single-cell and population-level techniques to perform a systematic study on the activity of human antimicrobial peptide LL37 on *E. coli* cells in exponential and stationary phases. In single-cell experiments, we quantified the time of AMP-induced cell death for *E. coli* cells and in the population-level experiments we measured the minimum bactericidal concentration (MBC) of LL37 peptides.

LL37 peptides do not specifically attack a cell-cycle-dependent pathway, but the action has a subtle, indirect dependence on cell age. In fact, it has been previously shown by Weisshaar lab that LL37 peptides preferentially bind around the septum of dividing cells. In this work, our single-cell observations confirm that cells at or near cell division stage are more susceptible to LL37 peptides. And since the frequency of cell division is reduced in the absence of nutrients, the action of LL37 peptides is slower on non-growing, starving *E. coli* cells as compared to that on exponentially growing cells.

Interestingly, this rate difference is not reflected in the MBC as we found that the MBC of LL37 peptides is lower in cultures of starving cells than in exponentially growing cultures. In this work we specifically used cultures of the same cell density to avoid inoculum effect, which has been demonstrated to be a significant factor for AMPs action [16, 24, 26].

The smaller MBC of LL37 in starving culture is counterintuitive as it is expected from an antibiotic with a higher rate of action. One possibility, which we hypothesize here, is that the action of LL37 peptides is triggered by accumulation of peptides on the membrane to a threshold, which is achieved at a higher MBC on exponentially growing cells since growth and surface expansion dilutes out the surface covered by LL37 peptides (Fig. 4B).

On the other hand, the surface of a non-growing cell remains in stagnation and LL37 peptides can accumulate over a long period of time, though at a lower rate, but can eventually reach a high surface concentration to trigger membrane rupture. A direct, quantitative study of this model remains to be investigated in future work.

The question we addressed in this work is of fundamental significance in the design of peptide antibiotics and their synthetic mimics. Bacteria in clinical infections are typically found in low nutrient or stationary phase conditions. Thus, this is crucial to design antibiotics agents that can efficiently attack the non-growing cells.

## Acknowledgments

This work was supported by the National Institute of Health grant R15-GM124640 and by the startup funds from California State University, Northridge.

## Footnotes

1 The death time is judged by the moment of peptide translocation in the target cells.

2 The tag is known to slightly modify the inhibitory concentration of peptides [24]

